## Supplementary File 1 for "Neural Representation of Time across Complementary Reference Frames"

**Supplementary File 1:** The reaction time of the corrected trials indicates that time is differently processed under internal and external perspectives[1]

| Fixed Effects[2] | df | F | p |
|---|---|---|---|
| Sequential Distance | 1, 6914 | 0.018 | 0.895 |
| Duration | 1, 6914 | 7.687 | 0.006 |
| Same vs. Different | 1, 6914 | 0.862 | 0.353 |
| Future vs. Past | 1, 6914 | 0.088 | 0.766 |
| Syllable Length | 1, 6914 | 33.222 | < 0.001 |
| Task Type | 1, 6914 | 5.845 | 0.016 |
| Task Type × Sequential Distance | 1, 6914 | 13.959 | < 0.001 |
| Task Type × Duration | 1, 6914 | 6.246 | 0.012 |
| Task Type × (Same vs. Different) | 1, 6914 | 70.490 | < 0.001 |
| Task Type × (Future vs. Past) | 1, 6914 | 0.002 | 0.965 |
| Task Type × Syllable Length | 1, 6914 | 0.961 | 0.327 |

[1] Linear Mixed Model Formula: RT ~ 1 + Task Type * (Sequential Distance + Duration + Same/Different + Future/Past + Syllable Length) + (1 | Participant)

[2] The significant effects were highlighted. We did not highlight the significant main effects if the corresponding interaction effects were also significant.
